# Intravenous midazolam alters short-interval paired-pulse TMS responses differently in younger and older adults

**DOI:** 10.64898/2026.06.20.733493

**Authors:** Keith M. McGregor, Seyed A. Safavynia, Thomas Novak, Ashton M. Weber, Jessica Wang, Joe R. Nocera, Anna Woodbury, Bruce Crosson, Paul S. García

## Abstract

**Objective:** Aging is associated with changes in cortical excitability and altered responsiveness to benzodiazepines, but the effects of benzodiazepine challenge on motor cortical paired-pulse physiology in older adults remain incompletely understood. We examined whether intravenous midazolam differentially modulates corticospinal excitability and short-interval paired-pulse transcranial magnetic stimulation (TMS) responses in younger and older adults.

**Methods:** Fifteen younger adults (18–35 years) and fifteen older adults (50–69 years) underwent single-pulse and paired-pulse TMS of the left primary motor cortex at baseline and during intravenous midazolam administration. Single-pulse motor evoked potential (MEP) amplitude was used to assess corticospinal excitability. Short-interval paired-pulse responses were quantified as the ratio of conditioned to unconditioned MEP amplitude.

**Results:** At baseline, younger adults showed greater corticospinal excitability than older adults, reflected by larger single-pulse MEP amplitudes (adjusted p = 0.04). Younger adults demonstrated paired-pulse inhibition at baseline, reflected by a conditioned/unconditioned MEP ratio below 1.0 (ratio = 0.73; adjusted p < 0.01), whereas older adults did not show inhibition and instead had a mean ratio above 1.0 (ratio = 1.25). Midazolam reduced single-pulse MEP amplitudes in both groups. During midazolam administration, paired-pulse inhibition was no longer observed in younger adults, and older adults continued to show no evidence of inhibition.

**Conclusions:** Younger and older adults differed in baseline corticospinal excitability and in short-interval paired-pulse TMS responses. Intravenous midazolam reduced corticospinal excitability and altered paired-pulse response patterns, eliminating baseline paired-pulse inhibition in younger adults while producing little measurable change in older adults. These findings suggest that aging may modify the net motor cortical response to benzodiazepine challenge. The results should be interpreted in relation to the paired-pulse stimulation parameters used and support further studies using complementary approaches to characterize age-related differences in inhibitory and facilitatory motor cortical circuits.

## 1. INTRODUCTION

Aging is associated with changes in central nervous system function, including alterations in cortical excitability, inhibitory signaling, and responsiveness to medications that act on γ-aminobutyric acid (GABA) pathways. GABA is the primary inhibitory neurotransmitter in the brain and contributes to the regulation of cortical excitability across the lifespan (Cuypers et al., 2021). Benzodiazepines act as positive allosteric modulators of the GABAA receptor and are widely used clinically for anxiolysis, sedation, and amnesia. However, benzodiazepine effects in older adults are often variable, with evidence of increased sensitivity, delayed clearance, and paradoxical excitation in some individuals (Albrecht et al., 1999; Sun et al., 2008). These age-related pharmacodynamic and behavioral differences raise important questions about how inhibitory physiology changes with aging and how such changes may be captured using noninvasive physiological measures.

Transcranial magnetic stimulation (TMS) provides a noninvasive approach for assessing motor cortical excitability and the balance of inhibitory and facilitatory circuit activity. Single-pulse TMS can be used to quantify corticospinal excitability through the amplitude of motor evoked potentials (MEPs). Paired-pulse TMS paradigms provide additional information about how a conditioning stimulus modifies the response to a subsequent test stimulus. One widely used paired-pulse paradigm is short-interval intracortical inhibition (SICI), in which a conditioning stimulus delivered shortly before a test stimulus reduces the resulting MEP amplitude (Kujirai et al., 1993). Short interval (2-3 ms interstimulus interval) paired pulse inhibition paradigms have commonly been interpreted as reflecting GABA_A_-associated inhibitory processes within primary motor cortex, although their magnitude is influenced by stimulation parameters, arousal, circadian state, baseline excitability, and other physiological factors (Cirillo et al., 2018; Lang et al., 2011; Ziemann et al., 2015). Thus, paired-pulse TMS responses are best understood as circuit-level physiological measures that can be sensitive to GABAergic modulation but are also shaped by the broader state of the motor system.

Studies examining age-related differences in paired-pulse inhibition have produced mixed results. Several reports suggest reduced paired pulse inhibition or altered inhibitory physiology in older adults (Heise et al., 2013; Hermans et al., 2018), whereas others have reported no age-related differences or more complex changes in corticospinal excitability across adulthood (Bhandari et al., 2016; Hehl et al., 2020). These inconsistencies may reflect differences in stimulation intensity, thresholding procedures, test MEP amplitude, task state, participant characteristics, or analytic approach. Age-related changes in GABA_A_ receptor density, subunit composition, synaptic organization, and cortical structure may also contribute to variability in inhibitory and facilitatory responses (Robinson et al., 2019; Rozycka and Liguz-Lecznar, 2017). As a result, age-related changes in paired-pulse TMS measures remain incompletely understood, particularly when assessed under pharmacological challenge.

Pharmacological manipulation provides a complementary approach for probing motor cortical physiology by examining how cortical excitability and paired-pulse responses change when inhibitory neurotransmission is modulated. In younger adults, benzodiazepines such as diazepam and lorazepam have been reported to increase SICI under conventional stimulation conditions, consistent with enhancement of GABA_A_-associated inhibitory signaling (Di Lazzaro et al., 2007; Premoli et al., 2017). However, responses to benzodiazepines may vary depending on the specific drug, dosing method, arousal state, baseline excitability, and stimulation parameters. Despite the widespread clinical use of benzodiazepines in older adults, relatively little is known about how intravenous benzodiazepine administration alters motor cortical paired-pulse responses in aging. This question is important because older adults may differ from younger adults not only in baseline inhibitory physiology, but also in how motor cortical circuits respond to pharmacological modulation.

In the present study, we examined age-related differences in corticospinal excitability and short-interval paired-pulse TMS responses before and during intravenous midazolam administration, a rapidly acting benzodiazepine that potentiates GABA_A_ receptor activity. Fifteen younger adults and fifteen older adults underwent TMS assessment of single-pulse MEP amplitude and paired-pulse MEP modulation at baseline and during midazolam infusion. We hypothesized that older adults would show lower baseline corticospinal excitability and reduced paired-pulse inhibition relative to younger adults. We further hypothesized that midazolam would alter corticospinal excitability and paired-pulse response patterns, with attenuated modulation in older adults. As paired-pulse responses reflect the net effect of inhibitory and facilitatory circuit recruitment under the stimulation parameters used, we interpreted the findings as evidence of age-related differences in motor cortical response to benzodiazepine challenge rather than as a direct receptor-specific assay of intracortical inhibition.

## 2. METHODS

### 2.1. Participants

Participants were recruited from a volunteer database and community postings, including younger (18–35 years) and older (50–69 years) adults. We did not include patients >70 years because of the possibility of adverse events associated with benzodiazepines in this demographic. Other inclusion criteria were: 1) sedentary lifestyle, defined as not engaging in structured physical activity and/or not accumulating ≥45 minutes of moderate physical activity most days of the week, 2) right-handedness, 3) native English speakers, and 4) approval by primary care physician for study participation. Exclusion criteria were: 1) history of depression or neurological disease (including Parkinson’s disease, Alzheimer’s disease, multiple sclerosis, stroke), 2) conditions contraindicated with TMS (e.g., seizure, tremor), 3) use of GABAergic medications (e.g., benzodiazepines, barbiturates, baclofen, gabapentin, other benzodiazepine agonists), 4) hospitalization within the past 6 months, 5) uncontrolled hypertension or diabetes, 6) inability to walk 400 meters, 7) significant cognitive executive impairment, defined as a Montreal Cognitive Assessment (MoCA) score <23 (Milani et al., 2018), 8) failure to provide informed consent.

Study personnel explained the purpose and potential risks of the experiment and obtained written informed consent from each participant following protocols approved by the Emory University Institutional Review Board (IRB) #IRB00079741 in compliance with the Helsinki Declaration. This study was reviewed and approved as human subjects research. The study was not registered as a clinical trial as it was designed as a mechanistic pharmacologic physiology study using TMS-derived neurophysiological outcomes. Participants were recruited between June 2017 and July 2021 from a volunteer database and community postings.

### 2.2. Study design

Participants were scheduled to complete three sessions of data acquisition, of which data for the current study were collected on Sessions 1 and 2 (**Figure 1**). In Session 1, following informed consent, participants underwent cognitive and behavioral screening and upper extremity motor testing (see *Assessments* section). In Session 2, participants underwent a motor thresholding protocol followed by two TMS assessments, one at baseline without drug (“No Drug” condition) and one in the presence of midazolam infusion (“Midazolam” condition).

**Figure 1.**
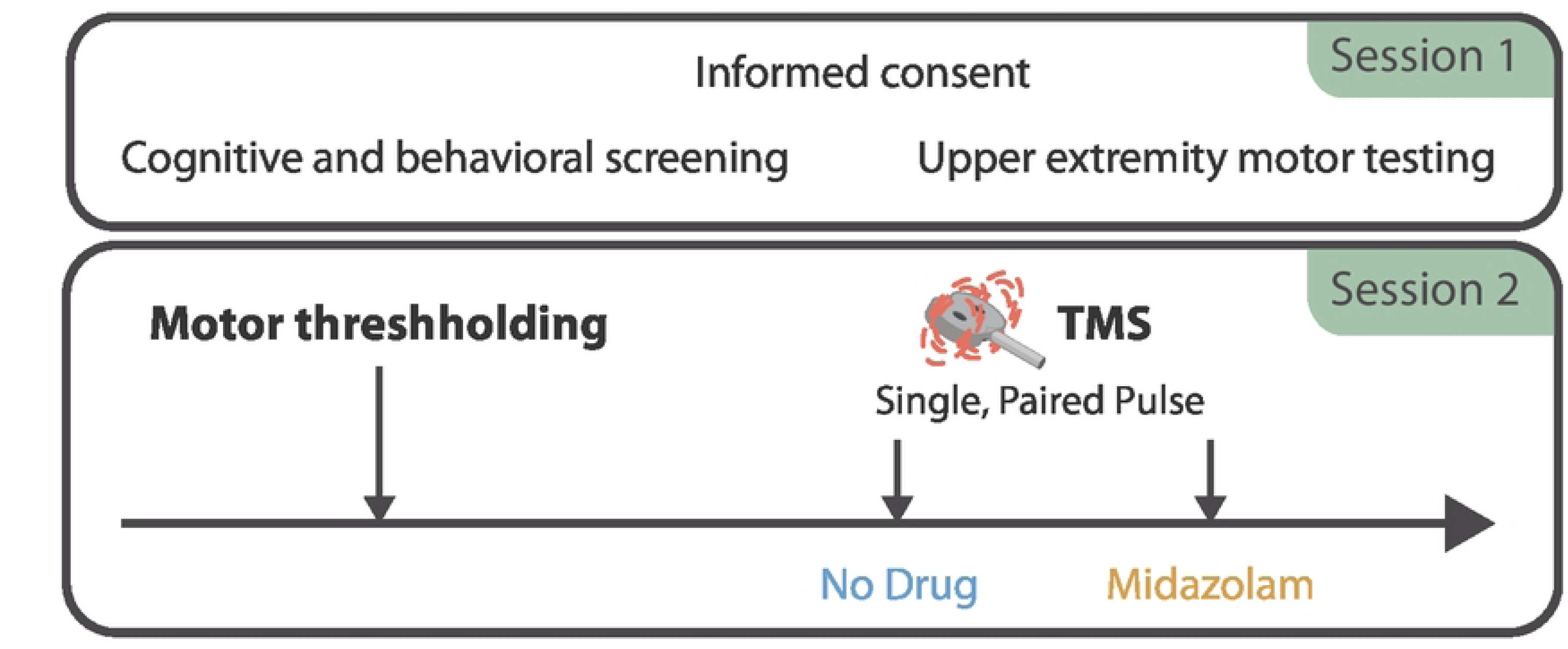
Study Flowchart. Participants underwent consent, behavioral screening and upper extremity motor testing on their initial visit (Session 1). On the second visit (Session 2), participants underwent motor thresholding followed by TMS assessment (see 2.4 TMS protocol) in no drug and midazolam treatment conditions (see 2.5 benzodiazepine protocol).

At the outset of Session 2, a physician placed an intravenous (IV) catheter in the participants’ upper extremity per standard clinical practice and flushed the catheter with saline (≤5 cc). Participants then transferred to a specialized chair (Magventure, Alpharetta, GA) for the TMS and benzodiazepine protocols. The headrest was removed to prevent participants from dozing. After confirming participant comfort level, the physician administered low-flow (2 L/min) oxygen to participants *via* a nasal sampling cannula connected to a gas analyzer. Standard anesthesia monitoring was performed (heart rate, blood pressure, pulse oximetry, end-tidal carbon dioxide sampling, respiratory rate, temperature). Heart rhythm was continually assessed *via* 5-lead electrocardiogram, and blood pressure was monitored at 5-minute intervals throughout the session.

### 2.3 Assessments

#### 2.3.1. Cognitive and behavioral screening

The MoCA was used to screen participants for cognitive impairment. The MoCA is a validated cognitive screening test with assessment domains spanning memory, visuospatial ability, executive function, attention, concentration, working memory, language, abstract reasoning, and orientation. Scores of ≥23/30 are considered cognitively normal given ethnic and cultural considerations (Milani et al., 2018). Other assessments (Timed Up-and-Go test, Halstead finger tapping task, Purdue pegboard (column and assembly), and nine-hole pegboard task) were used to screen for undiagnosed motor deficits. All motoric assessments were performed using the dominant hand. Assessment sessions did not exceed two hours to minimize participant fatigue.

### 2.3.1. Arousal

The participants’ level of arousal was intermittently assessed during pauses in TMS trials using the Observer’s Assessment of Alertness/Sedation Scale (OAA/S). The OAA/S has previously been used to determine the end of anesthesia emergence and the beginning of anesthesia recovery (Hesse et al., 2019). Briefly, participants unresponsive to repeated verbal stimuli combined with tactile non-painful stimulation receive an OAA/S score of <2. The lowest OAA/S score for each drug condition was recorded for each participant. All scores ranged between 3–5 (a score of 3 is an active verbal response to hearing one’s name loudly; a score of 5 is an appropriate verbal response with eye contact to verbal stimuli).

### 2.4. TMS protocol

#### 2.4.1. Electromyography (EMG)

EMG was continuously sampled from the first dorsal interosseous (FDI) muscle on both hands using BrainSight 2 (Rogue Research, Cambridge MA) with Ag/AgCl electrodes. Immediately prior to electrode placement, target sites were lightly abraded with alcohol wipes and a small quantity of conductive gel was applied. EMG recordings (trials) were triggered via transistor-to-transistor logic (TTL) pulse over a 250 ms window per trial (50 ms prestimulus window followed by 200 ms of response). A LabJack U3-LV analog to digital converter acquired amplified EMG traces with a 12-bit dual-channel analog input sampled at 3 kHz. Data were bandpass filtered from 10–1,000 Hz. Muscle activation was monitored during sessions with the oscilloscope software package integrated into the BrainSight neuronavigated positioning system.

#### 2.4.2. TMS acquisition

A MagVenture X100 magnetic stimulator with MagOption and B-60 60 cm butterfly coil (MagVenture, Alpharetta, GA) was used to stimulate the left primary motor cortex. The TMS acquisition began with an initial thresholding procedure targeting the right FDI muscle using a standard template brain set in Montreal Neurological Institute space. An 8x8 square grid was overlaid on the template brain with grid spacing at 1 cm. All stimulations were biphasic, oriented anterior-to-posterior. Stimulation and recording devices were synchronized using TTL pulses. The coil was placed tangential to the scalp with the handle pointing backward and 45° away from the midline for stimulation. The resting motor threshold (RMT) was defined as the scalp site (“hotspot”) corresponding to the lowest stimulator output sufficient to generate a magnetic evoked potential of at least 50 μV (peak-to-peak) in six of ten trials. Intertrial intervals varied from 4-7 seconds to minimize repetitive stimulation effects.

#### 2.4.3. Stimuli

Baseline (single-pulse) MEP measurements were obtained from ten stimulations of the left primary motor area FDI hotspot at 130% RMT. Short-interval paired-pulse motor cortical responses were assessed using a conditioning-test pulse TMS protocol. A conditioning TMS pulse set at 110% of RMT was applied to the left motor cortex FDI hotspot 3 milliseconds prior to administration of a “test pulse” at 143% of RMT to the left motor cortex. In conventional short-interval paired-pulse paradigms, the conditioning stimulus can reduce the test-pulse MEP amplitude, a response commonly interpreted as paired-pulse inhibition. Due to our interest in the variability of MEP response between age groups, we elected to employ paired-pulse inhibition using a fixed stimulation value relative to RMT instead of EMG-derived 1 mV peak-to-peak value. Ten paired-pulse MEP measurements were obtained for each participant; the inter-trial interval was varied between four and six seconds randomly to reduce anticipation of the next trial and mitigate repetitive stimulation effects.

### 2.5. Benzodiazepine protocol

Midazolam is a non-narcotic, anti-anxiety agent with low abuse potential (Schedule IV) according to Title 21 Code of Federal Regulations Part 1308.14. It is the most commonly administered benzodiazepine in clinical anesthesiology and is most often used for pre-procedural anxiolysis/sedation. Its effects are mainly mediated by its action of enhancing inhibitory current through the GABA_A_ receptor.

For the study, a board-certified anesthesiologist administered an intravenous (IV) midazolam bolus followed by IV midazolam infusion throughout the testing protocol. For participants <80 kg, 1 mg midazolam was administered as a bolus, followed by infusion of 2 mg midazolam/hour. Participants ≥80 kg received a 1.5 mg midazolam bolus followed by infusion of 3 mg midazolam/hour. Midazolam infusions were delivered *via* a Baxter AS50 infusion pump prepared with 1.0 mg midazolam in 20 cc normal saline for a final concentration of 0.05 mg/mL (participants <80 kg) or 1.5 mg midazolam in 20 cc normal saline for a final concentration of 0.075 mg/mL (participants ≥80 kg); the midazolam dosage was set at a fixed infusion rate of 40 cc/hr. The approximate plasma concentration target was 10–30 ng/mL. Following midazolam administration, we waited 3–5 minutes for the medication to circulate, while engaging in topical conversation with the participant. The actions of midazolam can be immediately reversed by intravenous administration of the benzodiazepine antagonist, flumazenil, which was readily available to the physician for all testing sessions.

### 2.6. Quantitative measures of MEP

MEP amplitudes were defined as the peak-to-peak amplitude of the motor response in the right FDI muscle (in µV) for 50 ms following cortical stimulation. MEP amplitudes were calculated for all individual trials over all stimuli (single-pulse, paired-pulse) and treatment (no drug, midazolam) conditions in all participants using custom MATLAB routines. We excluded individual trials with MEP amplitudes <50 µV as these trials likely represented a failure of TMS to elicit a motor response. For each stimulus and treatment condition, we included only participants with MEP amplitudes ≥50 µV in more than six of the ten conditions yielded n=15 younger and n=15 older participants for the no drug condition, and n=15 younger and n=12 older participants for the midazolam condition. To compare the effects of paired-pulse stimulation in the presence and absence of midazolam, we analyzed only data from participants with at least 6 remaining trials after exclusions across all conditions, yielding n=15 younger and n=11 older participants.

Motor evoked potential amplitudes were visually and programmatically screened for amplifier-induced clipping. The recording system had a maximum range of 1.56 mV due to hardware limitations. Trials in which peak-to-peak MEP values exceeded this threshold were marked as clipped and excluded from direct analysis. To preserve the continuity of the input–output (IO) curve and allow for more accurate modeling of excitability, these values were estimated using cubic spline interpolation (**Figure 2**). Interpolation was performed using the MATLAB function ‘interp1’ (MathWorks, Natick, MA) with ‘spline’ as the specified method. Clipped points were then replaced with interpolated estimates derived from this fit. Overall, 75 of 1,077 trials (7.0%) were interpolated; interpolated peak-to-peak amplitudes differed from raw traces by 7.9 ± 3.5% in younger subjects and 6.9 ± 4.4% in older subjects. A full categorization of interpolated trials and characteristics can be found in **Table S1**.

**Figure 2.**
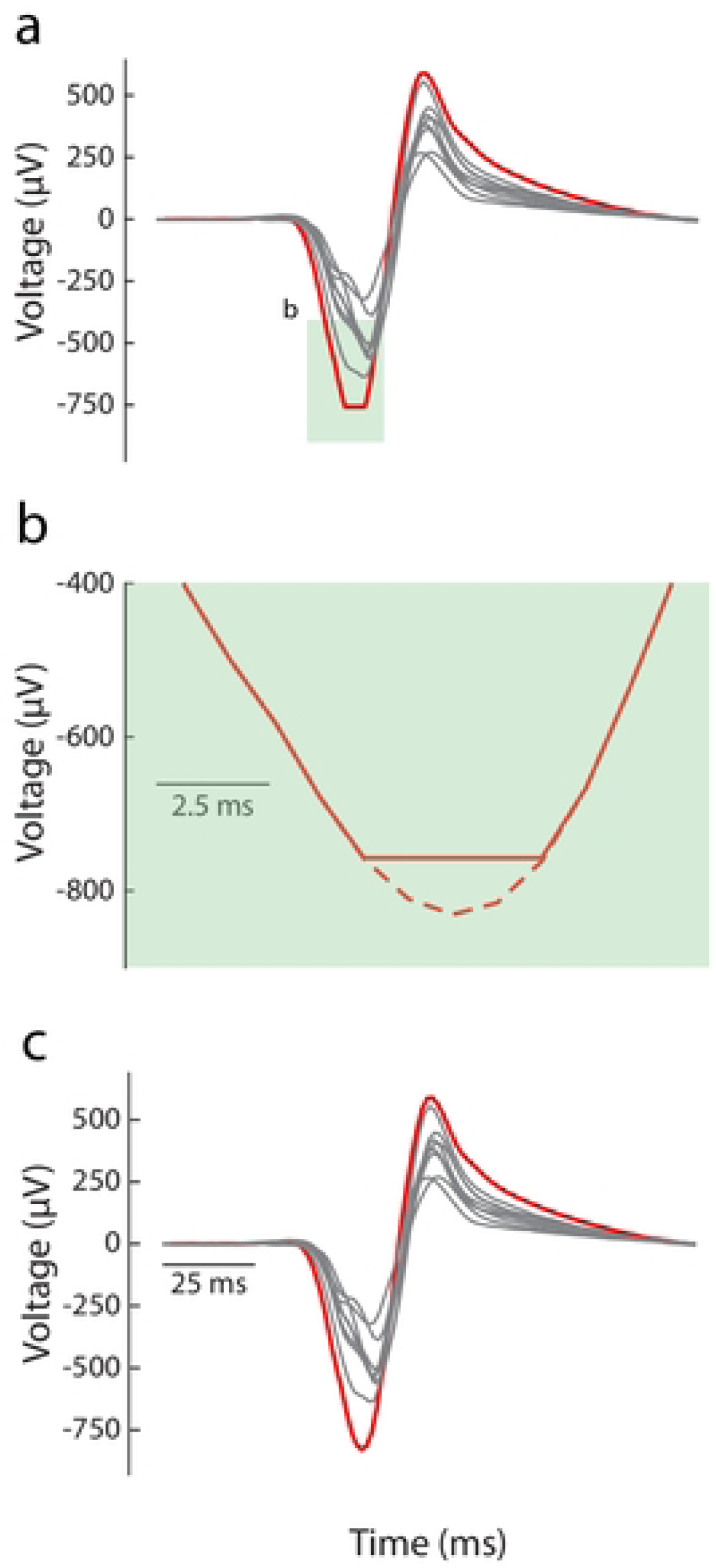
Interpolated MEPs. 2a: No drug single-pulse MEP traces for a representative participant. Unclipped traces are in gray, and clipped trace is in red. 2b: Magnification of clipped trace in the green shaded region of Figure 2a. Solid line represents raw data; dashed line represents interpolation calculation. 2c: interpolated trace.

### 2.7. Statistical comparisons

Demographic data were compared across groups using independent sample two-tailed t-tests at a significance of α = 0.05; contrasts are reported without correction. Average MEP amplitudes were calculated for all conditions in all participants. Normality of data was verified across all participants *via* the Shapiro-Wilk test. Single-pulse MEP amplitudes and paired-pulse inhibition ratios were compared across groups using independent sample t-tests at a significance level of α = 0.05. Changes in MEP amplitudes across conditions (single- *vs.* paired-pulse, no drug *vs.* midazolam) were compared within groups using paired t-tests at a significance level of α = 0.05. All statistical tests for MEPs were adjusted for multiple comparisons using Benjamini-Hochberg correction (Benjamini and Hochberg, 1995)); adjusted p-values are reported throughout the manuscript. Statistical comparisons were made in MATLAB (Mathworks, Danvers, MA) and JMP15 (SAS Institute, Cary, NC).

## 3. RESULTS

### 3.1. General characteristics

**Table 1** shows demographic data for participants. There were no significant differences in education. Older adults had a minimum MoCA score of 23, indicating intact cognitive function, considering cultural demographics. Motoric assessments demonstrated that all participants had upper extremity motor function within the age-adjusted range.

**Table 1.** Demographic data for younger (n = 16) and older (n = 16) participants.

|  | Younger Adults | Older Adults | t(df) | p-value |
| --- | --- | --- | --- | --- |
| **N** | 16 (4 Females) | 16 (5 Females) |  |  |
| **Age (yr)** | 24.3 (5.2) | 60.8 (6.3) | t(30) = 14.6 | **< 0.001\*** |
| **Weight (kg)** | 87.2 (15.5) | 83.8 (19.8) | t(30) = .56 | 0.57 |
| **Height (cm)** | 179.1 (9.7) | 170.3 (9.4) | t(30) = 2.7 | **0.01\*** |
| **Education** | 14.9 (3.6) | 14.8 (3.9) | t(30) = .07 | 0.94 |
| **MoCA** | 28.2 (1.4) | 26.9 (2.5) | t(26) = 1.8 | 0.07 |

### 3.2. Pulse MEP amplitude

Consistent with previous studies in the literature, in the absence of drug, older participants had smaller single-pulse MEP amplitudes compared to younger participants (**Figure 3**). Individual (grey) and average (black) single-pulse MEP traces are shown for representative younger and older participants (**Figure 3a**). Across subjects, younger adults had higher baseline MEP amplitudes (mean ± standard deviation: 788.3 ± 389.9 µV) compared to older adults (509.5 ± 298.5 µV, adjusted p= 0.04) (**Figure 3c**).

**Figure 3.**
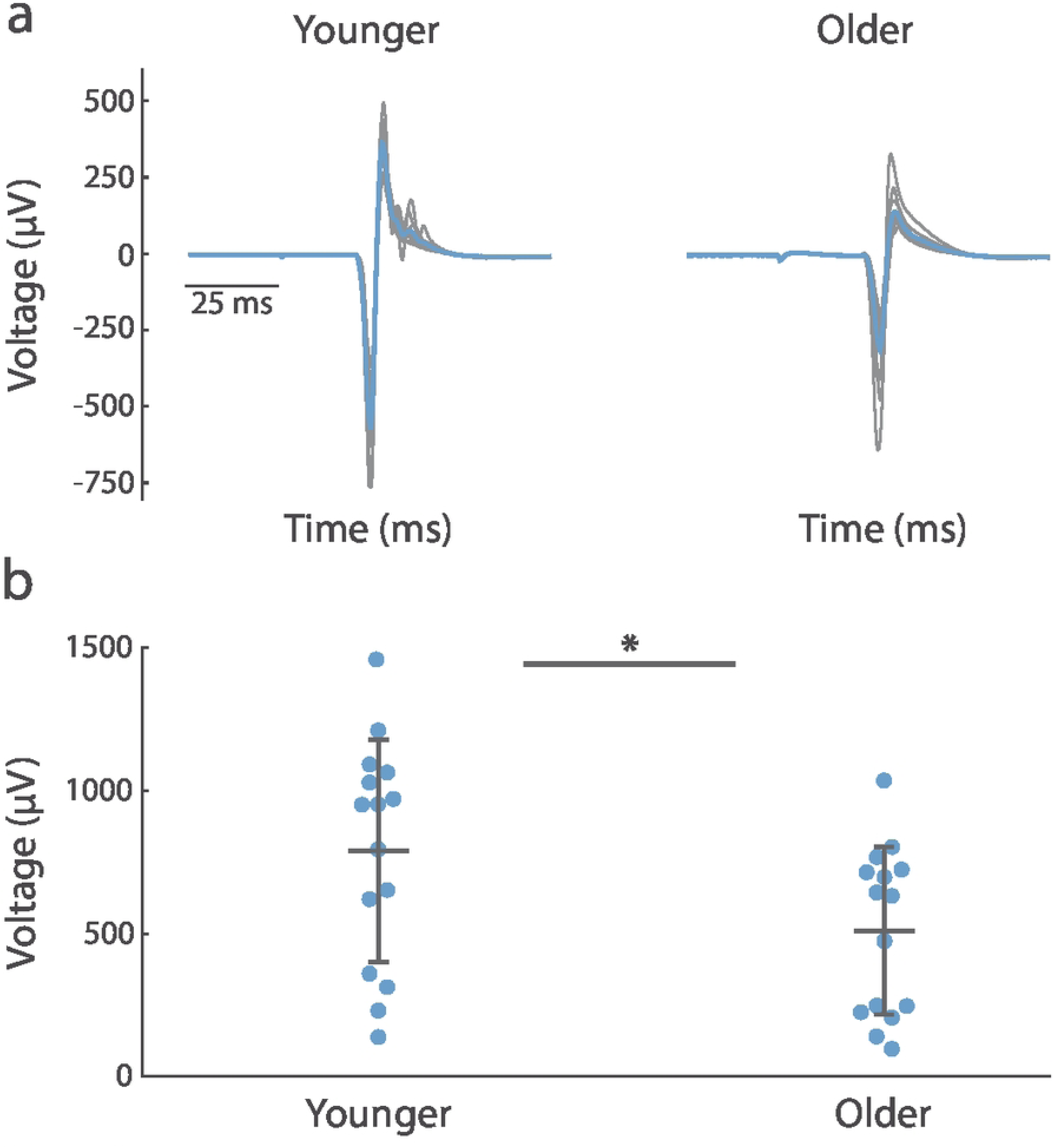
Single-pulse MEP amplitudes are smaller in older versus younger participants. 3a: Average MEP (blue) and individual trial (gray) single-pulse MEPs in no drug condition for a representative younger (left) and older (right) participant. 3b: Beeswarm plot of average MEP amplitudes for younger (n=15) versus older (n=15) participants. Cross represents mean (horizontal line) and standard deviation (vertical line with anchors). *adjusted p<0.05.

### 3.3. Effects of age on paired-pulse MEP amplitude and paired-pulse inhibition ratio

In the absence of drug, there was a significant difference in MEP amplitude across age groups (**Figure 4**). 13 of 15 younger participants had decreased MEP amplitudes for paired-pulse (PP) compared to single-pulse (SP) stimulation (PP 546.6 ± 274.0 µV vs. SP 788.3 ± 389.9 µV, adjusted p=0.0089) (**Figure 4a**). By contrast, only 6 of 15 older participants had decreased MEP amplitudes for paired-pulse compared to single-pulse stimulation (PP 605.5 ± 400.8 µV vs. SP 509.5 ± 298.5 µV, adjusted p=0.17).

**Figure 4.**
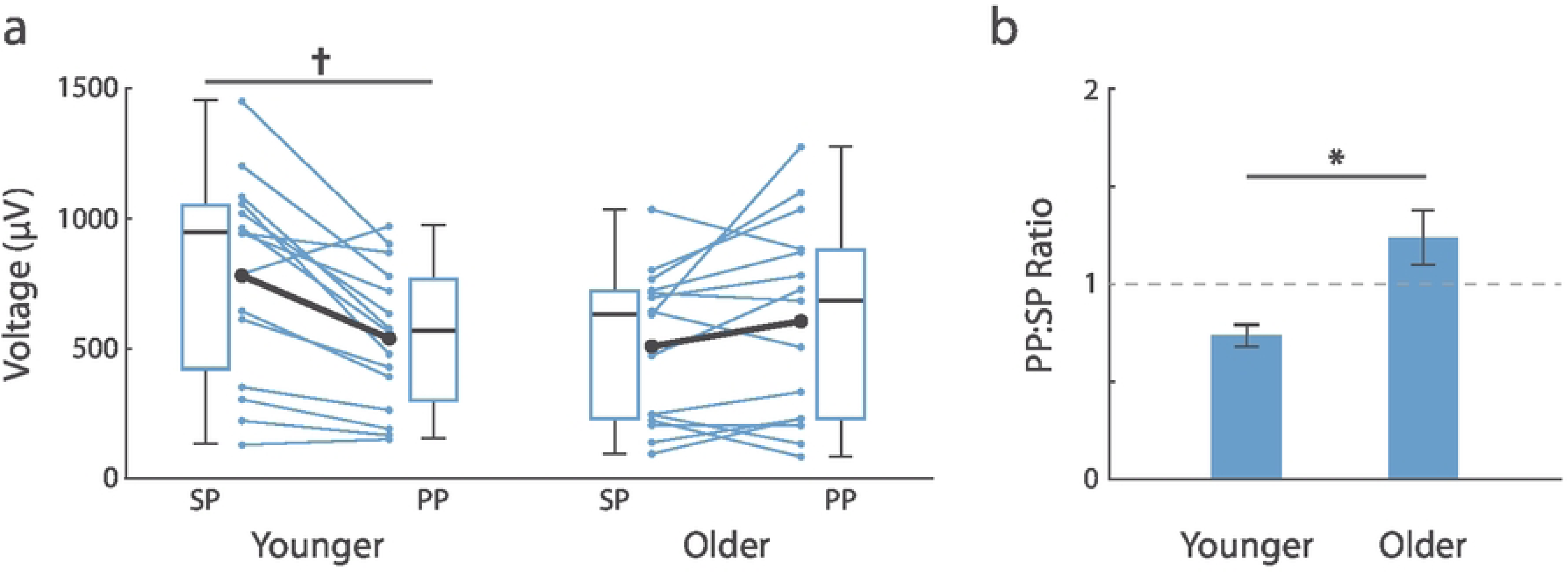
Age-related MEP amplitude modulation in response to paired-pulse stimulation. 4a: Average MEP amplitudes for each subject (blue paired dots) for single-pulse (SP) and paired-pulse (PP) stimuli in no drug condition for younger (n = 15) and older (n = 15) participants. Large black paired dots indicate average group MEP amplitude. Population data represented as box-and-whisker plots. For boxplots, black horizontal line is median, box is interquartile range, anchors are range. 4b: Ratio (mean ± standard error) of MEP amplitudes in paired-pulse compared to single-pulse stimuli for younger and older participants (paired-pulse ratio). ^†^adjusted p<0.01.

In the absence of drug, paired-pulse inhibitionwas only present in younger adults (**Figure 4b**). Younger adults demonstrated a paired-pulse to single-pulse ratio of 0.73 (95% confidence interval (CI) [0.61 0.85]). Older adults did not demonstrate any effect of paired-pulse inhibition (ratio 1.25, 95% CI [0.95 1.55]). The difference in paired-pulse inhibition ratios was significant across age groups (adjusted p=0.015).

### 3.4. Effects of midazolam infusion on unconditioned (single-pulse) MEP amplitude

Younger and older participants exhibited decreased MEP amplitude in the presence of midazolam (**Figure 5**). Mean ± standard deviation MEP traces are shown for a representative older and younger participant in **Figure 5a** in no drug (blue) and midazolam (orange) conditions. MEPs were decreased in the presence of midazolam in the majority of younger (11/15) and older (9/12) participants (**Figure 5b**, paired dots). The reduction in MEP amplitude following midazolam administration was significant in younger adults (no drug: 788.3 ± 389.9 µV *vs.* midazolam: 565.3 ± 350.1 µV; adjusted p = 0.04) but not older adults (no drug: 554.5 ± 291.0 µV *vs.* midazolam: 397.0 ± 421.8 µV, adjusted p = 0.17).

**Figure 5.**
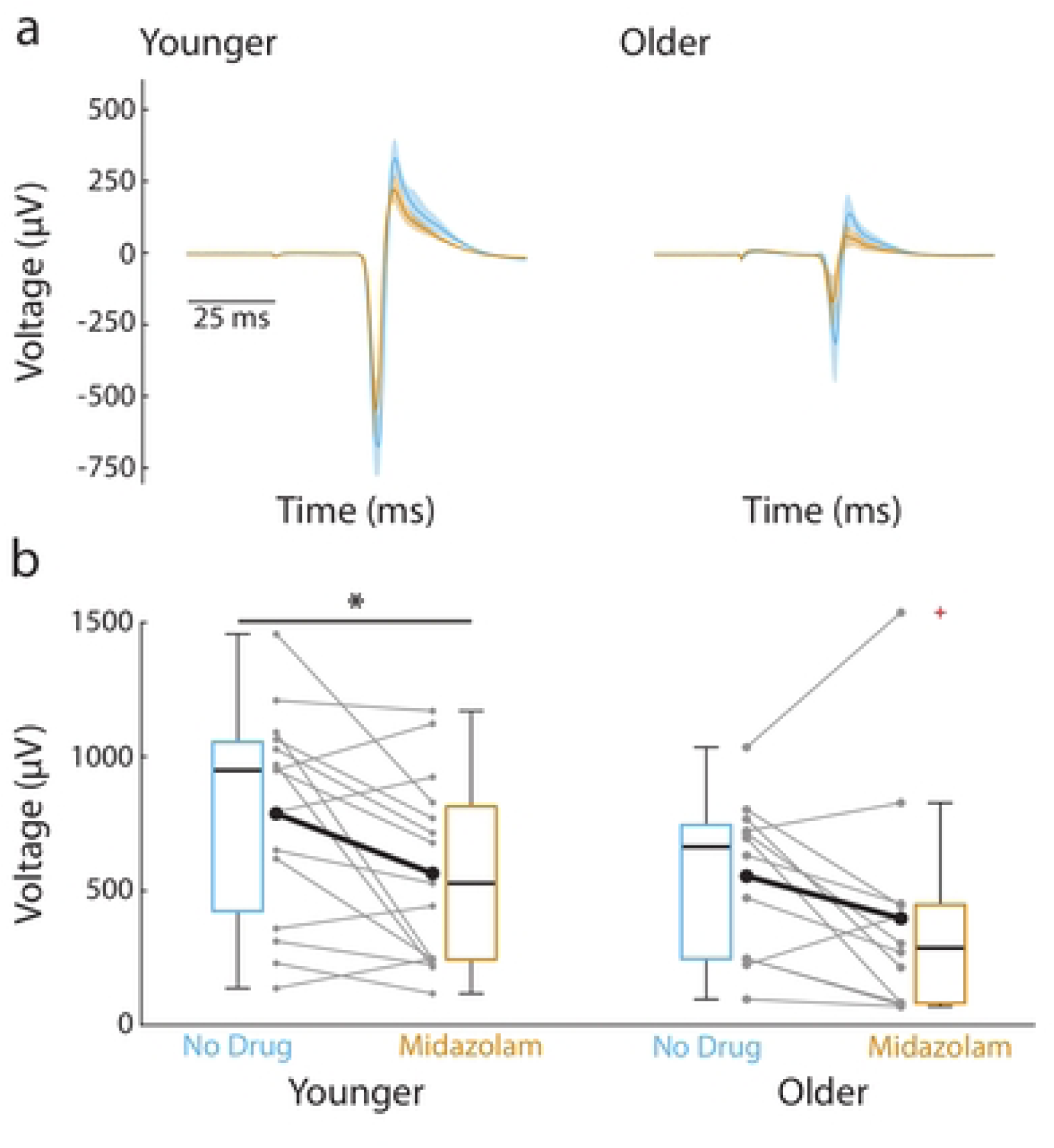
Midazolam reduces MEP amplitude in younger and older participants. 5a: Average MEP (solid lines) and standard deviations (shaded regions) in no drug and midazolam (Midaz) treatment conditions for a representative younger (left) and older (right) subject. 5b: Individual subject MEP amplitudes (gray paired dots) for single-pulse (SP) stimuli in no drug and midazolam treatment conditions for younger (n = 15) and older (n = 12) participants. Large black paired dots represent average group MEP amplitude. Population data represented as box-and-whisker plots for no drug (blue) and midazolam (orange) treatment conditions; black line is median, box is interquartile range, anchors are range, + is outlier. *adjusted p<0.05.

### 3.5. Effects of midazolam infusion on paired-pulse MEP amplitude and paired-pulse inhibition ratio with respect to age

In the presence of midazolam, there was no difference in MEP amplitudes for paired-pulse compared to single-pulse stimulation in both younger (PP 580.5 ± 336.0 µV vs. SP 565.3 ± 350.1 µV, adjusted p=0.77) and older adults (PP 476.7 ± 364.1 µV vs. SP 397.0 ± 421.8 µV, adjusted p=0.38; **Figure 6a**). Further, there was no difference in paired-pulse MEP amplitudes in baseline versus midazolam conditions for younger (adjusted p = 0.57) or older (adjusted p = 0.052) adults.

**Figure 6.**
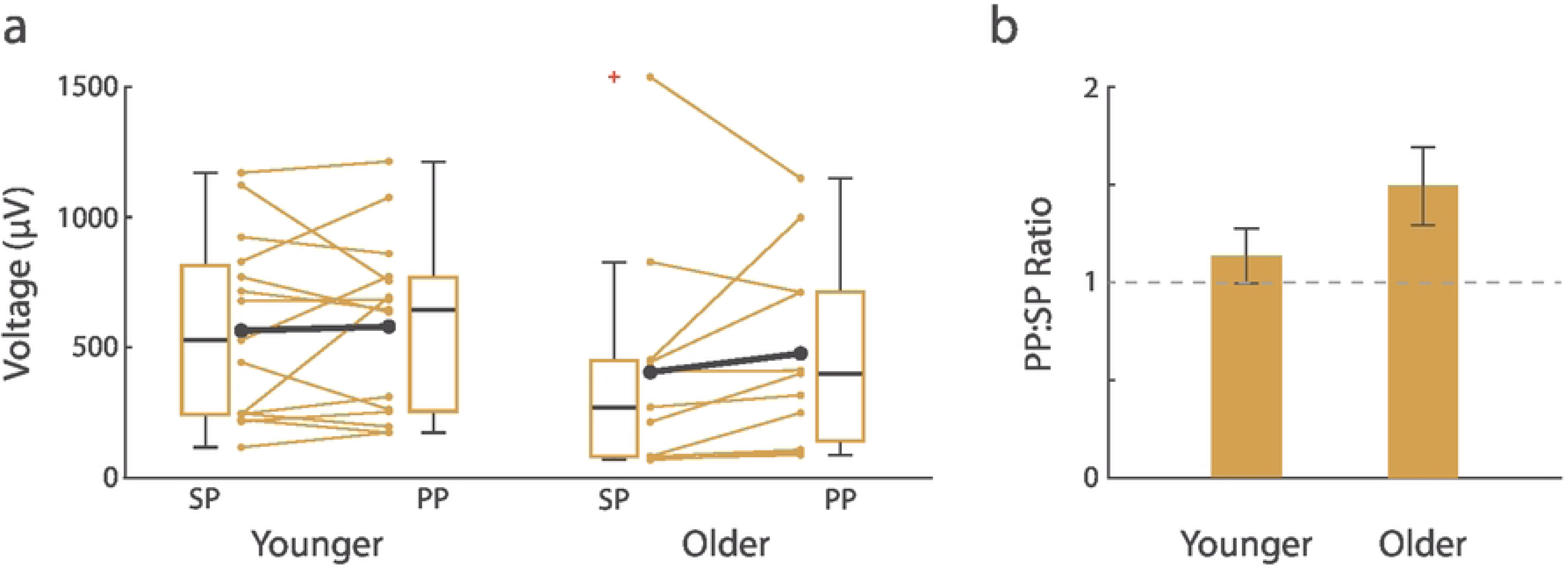
Paired pulse inhibition is abolished in the presence of midazolam. 6a: Average MEP amplitudes for each subject (colored paired dots) for single-pulse (SP) and paired-pulse (PP) stimuli in midazolam condition for younger (n = 15) and older (n = 11) participants. Large black paired dots indicate average group MEP amplitude. Population data represented as box-and-whisker plots. For boxplots, the black horizontal line is the median, the box is the interquartile range, anchors are range, + is outlier. 6b: Ratio (mean ± standard error) of MEP amplitudes in paired-pulse compared to single-pulse stimuli for younger and older participants (paired-pulse ratio).

In the presence of midazolam, paired-pulse inhibition was not present in younger or older adults (**Figure 6b**). Younger adults had a paired-pulse inhibition ratio of 1.14 (95% CI [0.83 1.45]); older adults had a paired-pulse inhibition ratio of 1.62 (95% CI [1.15 2.09]). There were no differences in paired-pulse inhibition ratios across age groups (adjusted p=0.19).

## 4. DISCUSSION

The present study examined age-related differences in corticospinal excitability and short-interval paired-pulse TMS responses before and during intravenous midazolam administration. At baseline, younger adults exhibited greater corticospinal excitability and clear paired-pulse inhibition, whereas older adults showed lower single-pulse MEP amplitudes and no consistent paired-pulse inhibition. Following midazolam administration, single-pulse MEP amplitudes were reduced, paired-pulse inhibition was no longer observed in younger adults, and older adults continued to show absent or facilitatory paired-pulse responses. These findings suggest that age and benzodiazepine challenge alter the net motor cortical response to paired-pulse stimulation. Rather than indicating a uniform enhancement of inhibition following pharmacologic potentiation of GABA_A_ receptors, the results suggest that paired-pulse TMS responses reflect the combined influence of baseline excitability, stimulation parameters, drug state, and age-related differences in inhibitory and facilitatory circuit recruitment.

To our knowledge, this is among the first studies to examine age-related differences in motor cortical paired-pulse responses during intravenous benzodiazepine administration. The findings are relevant to aging neurophysiology because benzodiazepines are commonly used in clinical settings and older adults often show variable pharmacodynamic and behavioral responses to these medications. The present results suggest that TMS-derived measures of cortical excitability and paired-pulse modulation may provide useful systems-level information about age-related differences in motor cortical response to pharmacologic challenge, while also underscoring the need for cautious interpretation of paired-pulse measures as receptor-specific indices.

### 4.1. MEP amplitude and variability with aging

Despite matched stimulation parameters based on individual motor thresholds, younger adults exhibited higher single-pulse MEP amplitudes than older adults, consistent with prior evidence that aging is associated with changes in motor cortical excitability (Bhandari et al., 2016). Reduced MEP amplitudes in older adults may reflect multiple factors, including changes in cortical excitability, alterations in inhibitory and excitatory neurotransmission, cortical atrophy, differences in coil-to-cortex distance, or age-related changes in peripheral motor output (Cuypers et al., 2020; Janssen et al., 2000; Porges et al., 2021; Salat et al., 2004). Although stimulation was scaled to individual resting motor threshold, threshold-based normalization does not fully eliminate the influence of anatomical and physiological variability across age groups.

The reduction in MEP amplitude during midazolam administration further indicates that the drug altered corticospinal excitability. This effect was most evident in younger adults but was directionally similar in older adults. Because MEP amplitude is influenced by cortical, spinal, and peripheral factors, the observed reduction should not be interpreted as a direct measure of GABA_A_ receptor function. Instead, it suggests that intravenous midazolam altered the overall excitability of the corticospinal system under the present stimulation conditions.

### 4.2. Short-interval paired-pulse responses with age

In the drug-naïve condition, younger adults showed consistent paired-pulse inhibition, whereas older adults showed more heterogeneous responses, including facilitation in several participants. This finding is broadly consistent with prior work suggesting that aging may alter inhibitory physiology within the motor system, although the direction and magnitude of age effects have varied across studies (Bhandari et al., 2016; Heise et al., 2013; Hermans et al., 2018). Such variability may reflect differences in stimulation parameters, baseline MEP amplitude, thresholding approach, task state, or participant characteristics.

The presence of facilitation rather than inhibition in several older adults may indicate a shift in the balance of inhibitory and facilitatory circuit recruitment. Age-related changes in GABA_A_ receptor density, subunit composition, synaptic organization, and cortical structure could contribute to such differences (Caspary et al., 1999; Pandya et al., 2019; Pirker et al., 2000; Robinson et al., 2019; Rozycka and Liguz-Lecznar, 2017). However, the current data do not isolate a specific receptor-level mechanism. The lower baseline MEP amplitudes in older adults also raise the possibility that ratio-based paired-pulse measures were influenced by reduced dynamic range or floor effects. Adaptive or threshold-tracking paradigms that adjust stimulation intensity to maintain a consistent MEP output may better characterize paired-pulse dynamics in older adults (Cirillo et al., 2018; Mooney et al., 2018).

### 4.3. Short-interval paired-pulse responses with midazolam

Contrary to our expectation, midazolam did not enhance paired-pulse inhibition in either age group. In younger adults, paired-pulse inhibition observed at baseline was no longer present during midazolam administration. In older adults, paired-pulse inhibition remained absent and responses were frequently facilitatory. These findings contrast with prior pharmacological TMS studies reporting increased SICI following oral benzodiazepines such as diazepam, lorazepam, or alprazolam (Di Lazzaro et al., 2007; Ferland et al., 2021; Florian et al., 2008; Teo et al., 2009). Several factors may account for this difference, including drug-specific pharmacology, route of administration, arousal state, baseline excitability, and stimulation parameters.

The present findings do not refute the relationship between conventional SICI and GABA_A_-associated inhibitory physiology. Rather, they indicate that short-interval paired-pulse responses under the present experimental conditions should be interpreted as net physiological responses of interacting inhibitory and facilitatory circuits. Midazolam may reduce overall corticospinal excitability while also changing the balance of circuit recruitment evoked by paired-pulse stimulation. At a microcircuit level, paired-pulse responses likely reflect the ensemble activity of multiple interneuron populations across cortical layers, including fast-spiking parvalbumin-positive interneurons and other inhibitory subtypes that differentially regulate cortical output (Tremblay et al., 2016). Age-related changes in receptor composition or pharmacologic sensitivity may contribute to the observed pattern, but these mechanisms cannot be determined directly from the current TMS measures.

The route of administration may also be relevant. Most prior pharmacological TMS studies of benzodiazepines have used oral agents, which have slower onset and different pharmacokinetic profiles than intravenous midazolam. Intravenous administration produces a more rapid drug effect, but plasma concentrations were not measured in this study, and central nervous system exposure may still vary across individuals. Age-related differences in benzodiazepine pharmacokinetics and pharmacodynamics could therefore contribute to variability in paired-pulse responses, particularly in the older group. Collectively, the results suggest that benzodiazepine challenge does not uniformly enhance paired-pulse inhibition and that aging may alter the motor cortical response to pharmacologic modulation.

### 4.4. Limitations

Several limitations should be noted. First, the sample size was modest, particularly after exclusions in the midazolam condition, which reduced power to detect age-by-drug interactions and limited characterization of inter-individual variability. Second, the paired-pulse protocol used a conditioning stimulus at 110% of resting motor threshold rather than a classical subthreshold SICI paradigm. Because paired-pulse responses are sensitive to stimulus intensity, this protocol may have recruited a different balance of inhibitory and facilitatory circuits and should be interpreted as reflecting this specific stimulation configuration rather than canonical SICI alone (Kujirai et al., 1993). Third, stimulation intensities were fixed relative to resting motor threshold rather than adjusted to produce a fixed MEP output. Given smaller baseline MEPs in older adults and further MEP reductions with midazolam, ratio-based measures may have been influenced by reduced dynamic range or floor effects, particularly in the older group (Cirillo et al., 2018; Mooney et al., 2018).

Fourth, although midazolam was administered under controlled anesthetic monitoring, plasma drug concentrations were not measured, limiting certainty regarding equivalent drug exposure across participants. Fifth, arousal was assessed intermittently with the OAA/S rather than continuously with electrophysiologic measures, leaving open the possibility that state-related fluctuations contributed to variability in paired-pulse responses. Sixth, although coil placement was standardized to the motor hotspot, the absence of subject-specific MRI-guided neuronavigation may have introduced spatial variability in stimulation, particularly given age-related anatomical differences such as increased coil-to-cortex distance. Finally, the older group was limited to healthy adults aged 50–69 years, which constrains generalizability to adults over age 70 and to clinical populations.

More broadly, paired-pulse TMS measures are associated with inhibitory physiology but reflect the net output of interacting cortical, corticospinal, and potentially spinal processes rather than a direct receptor-specific assay. Future studies using canonical subthreshold SICI, threshold-tracking methods, individualized MRI-guided neuronavigation, direct pharmacokinetic sampling, continuous arousal monitoring, and multimodal imaging will be important for clarifying the mechanisms underlying age-related differences in pharmacologic modulation of motor cortical physiology.

## 5. CONCLUSION

In summary, younger and older adults differed in baseline corticospinal excitability and short-interval paired-pulse TMS responses. Younger adults showed paired-pulse inhibition at baseline, whereas older adults did not. Intravenous midazolam reduced corticospinal excitability and eliminated baseline paired-pulse inhibition in younger adults, while producing little measurable change in paired-pulse response patterns in older adults. These findings suggest that aging may modify the net motor cortical response to benzodiazepine challenge. The results should be interpreted in relation to the paired-pulse stimulation parameters used and support further work examining how age, pharmacologic state, and inhibitory and facilitatory circuit recruitment interact to shape TMS measures of motor cortical physiology.

## Data Availability Statement

The de-identified data underlying the results reported in this article, along with analysis code and a data dictionary, will be made available in a public repository prior to publication.

## Acknowledgements

The views expressed in this work do not necessarily reflect those of the Department of Veterans Affairs or the United States Government. The funders had no role in study design, data collection and analysis, decision to publish, or preparation of the manuscript. The de-identified minimal dataset underlying the results reported in this article, along with analysis code, a data dictionary, and supporting documentation, is available in Zenodo at https://doi.org/10.5281/zenodo.20734853. The authors thank Holly Hudson, Lisa Calas, Dr. Kevin A. George, Bryana Whitaker Hardin, Dr. James Snitzer, Tuan Cassim, Dr. Lisa C. Krishnamurthy and Kevin Mammino for their contributions to this project. We would also like to thank Dr. Venkatagiri Krishnamurthy and Dr. Jeffrey Boatright for valuable discussions.

Conceptualization: Keith M. McGregor, Paul S. García, Bruce Crosson, Joe R. Nocera. Methodology: Keith M. McGregor, Seyed A. Safavynia, Paul S. García, Bruce Crosson, Joe R. Nocera.

Investigation: Keith M. McGregor, Paul S. García, Anna Woodbury.

Data curation: Keith M. McGregor, Seyed A. Safavynia, Ashton M. Weber, Jessica Wang, Thomas Novak.

Formal analysis: Keith M. McGregor, Seyed A. Safavynia, Ashton M. Weber, Jessica Wang, Thomas Novak.

Funding acquisition: Bruce Crosson, Keith M. McGregor, Paul S. García, Joe R. Nocera.

Project administration: Keith M. McGregor, Paul S. García.

Supervision: Keith M. McGregor, Paul S. García, Bruce Crosson, Joe R. Nocera.

Writing – original draft: Keith M. McGregor, Seyed A. Safavynia, Paul S. García, Ashton M. Weber, Thomas Novak, Jessica Wang.

Writing – review & editing: Keith M. McGregor, Seyed A. Safavynia, Thomas Novak, Ashton M. Weber, Jessica Wang, Joe R. Nocera, Anna Woodbury, Bruce Crosson, Paul S. García.

---

**Table S1.** Characteristics of MEP trials with interpolated peak-to-peak amplitudes.

|  |  | Younger |  |  | Older |  |  |
| --- | --- | --- | --- | --- | --- | --- | --- |
|  |  | No Drug | Midaz | Total | No Drug | Midaz | Total |
| SP | # Trials (%) | 27/141 (19.2) | 8/140 (5.7) | 35/281 (12.5) | 2/151 (1.3) | 10/105 (9.5) | 12/256 (4.7) |
| | Raw MEP, $\mu\text{V}$<br>(mean (SD)) | 1,275.7 (98.0) | 1,297.8 (90.1) | 1,280.8 (95.4) | 1,197.9 (20.9) | 1,433.8 (51.1) | 1,394.5 (103.0) |
| | Interpolated MEP, $\mu\text{V}$<br>(mean (SD)) | 1,374.0 (120.2) | 1,419.1 (113.0) | 1,384.3 (118.5) | 1,254.1 (32.5) | 1,536.2 (114.5) | 1,489.2 (151.3) |
| | %$\Delta$ (SD) | 7.7 (3.8) | 9.3 (2.2) | 8.0 (3.5) | 4.7 (4.5) | 7.0 (4.8) | 6.7 (4.6) |
| PP | # Trials (%) | 2/137 (1.5) | 8/137 (5.8) | 10/274 (3.7) | 14/146 (9.6) | 4/120 (3.3) | 18/266 (6.8) |
| | Raw MEP, $\mu\text{V}$<br>(mean (SD)) | 1,359.2 (39.4) | 1,343.0 (83.0) | 1,346.3 (74.7) | 1,230.8 (73.9) | 1,324.1 (137.9) | 1,251.5 (95.5) |
| | Interpolated MEP, $\mu\text{V}$<br>(mean (SD)) | 1,439.5 (86.0) | 1,450.6 (89.5) | 1,448.3 (84.1) | 1,323.4 (117.3) | 1,406.6 (152.2) | 1,341.9 (126.0) |
| | %$\Delta$ (SD) | 5.9 (3.3) | 8.1 (3.8) | 7.6 (3.6) | 7.4 (4.6) | 6.2 (3.4) | 7.1 (4.3) |
| Total | # Trials (%) | 29/278 (10.4) | 16/277 (5.8) | 45/555 (8.1) | 16/297 (5.4) | 14/225 (6.2) | 30/522 (5.8) |
| | Raw MEP, $\mu\text{V}$<br>(mean (SD)) | 1,281.5 (97.2) | 1,320.4 (86.9) | 1,295.3 (94.5) | 1,226.7 (69.9) | 1,402.4 (94.0) | 1,308.7 (120.2) |
| | Interpolated MEP, $\mu\text{V}$<br>(mean (SD)) | 1,378.5 (118.2) | 1,434.8 (99.8) | 1,398.5 (114.1) | 1,314.7 (112.0) | 1,499.2 (134.6) | 1,400.8 (152.9) |
| | %$\Delta$ (SD) | 7.5 (3.7) | 8.7 (3.0) | 7.9 (3.5) | 7.1 (4.5) | 6.8 (4.3) | 6.9 (4.4) |

Abbreviations: MEP, motor-evoked potential; PP, paired-pulse; SD, standard deviation; SP, single-pulse; % $\Delta$ , percent change; $\mu\text{V}$ , microvolts

